# Individual slow wave morphology is a marker of ageing

**DOI:** 10.1101/374397

**Authors:** Péter P Ujma, Péter Simor, Axel Steiger, Martin Dresler, Róbert Bódizs

## Abstract

Slow wave activity is a hallmark of deep NREM sleep. Scalp slow wave morphology is stereotypical, it is highly correlated with the synchronized onset and cessation of cortical neuronal firing measured from the surface or depth of the cortex, strongly affected by ageing, and these changes are causally associated with age-related cognitive decline. We investigated how normal ageing affects the individual morphology of the slow wave, and whether these changes are captured by the summary slow wave parameters generally used in the literature. We recorded full-night polysomnography in 159 subjects (age 17-69 years) and automatically detected slow waves using six different detection methods to ensure methodological robustness. We established individual slow morphologies at 501 data points for each subject and also calculated the individual average slow wave amplitude, average ascending and descending slope steepness and the total number of slow waves (gross parameters). Using LASSO penalized regression we found that fine-grained slow wave morphology is associated with age beyond gross parameters, with young subjects having faster slow wave polarity reversals, suggesting a more efficient initiation and termination of slow wave down- and upstates. Our results demonstrate the superiority of the high-resolution slow wave morphology as a biomarker of ageing, and highlights state transitions as promising targets of restorative stimulation-based interventions.

## Introduction

Electroencephalographic recordings in NREM sleep are characterized by slow, large amplitude waves (“slow waves”). These slow waves are cortical in origin (Steriade et al., 1993; Amzica and Steriade, 1995; Csercsa et al., 2010), with thalamic regulation (Crunelli and Hughes, 2010; David et al., 2013) and they reflect the rhythmic, highly synchronized onset and cessation of cortical neuron firing (Steriade et al., 1993; Nir et al., 2011). This particular pattern of neural firing has an important role in what is perhaps the most important function of NREM sleep: the normalization of synaptic connections formed during previous wakefulness (Tononi and Cirelli, 2014). In line with this observation, slow waves are more frequent and have greater amplitude in subjects and experimental conditions where stronger synaptic connections are expected to form before sleep: in younger subjects, whose synaptic plasticity is greater in general (Carrier et al., 2001; Carrier et al., 2011; Kurth et al., 2012; Feinberg and Campbell, 2013; Pótári et al., 2017) or in case of sleep deprivation (Borbely et al., 1981) or intensive, enriched wakefulness (Huber et al., 2004).

Slow waves – even when recorded from the scalp – accurately reflect the temporal dynamics of the synchronized onset and cessation of neural firing, as well as its spatial extent over the cortical mantle below the scalp electrodes (Nir et al., 2011). The detailed morphology of slow waves (slope steepness, minor structural differences, frequency) reflects the characteristics of neural firing, and it changes as a function of age and previous wakefulness: steeper, larger, smoother slow waves indicate stronger synchronized firing and therefore stronger synaptic connections (Huber et al., 2007; Riedner et al., 2007). As it is expected, this pattern is more common in young subjects, early in the night and after extended or more active wakefulness (Vyazovskiy et al., 2008; Bersagliere and Achermann, 2010; Colrain et al., 2010; Huber et al., 2013).

The age-related reduction of slow wave activity is well-known (Carrier et al., 2001; Carrier et al., 2011). Slow wave density and amplitude is lower in older individuals and it is especially associated with cognitive decline (Dresler et al., 2014; Yaffe et al., 2014), while retained slow wave activity at a higher age is associated with better cognitive and physical health (Anderson and Horne, 2003; Mazzotti et al., 2014). Our recent study (Pótári et al., 2017) has shown that high IQ is associated with a significantly attenuated age-related reduction in slow wave spectral power, indicating that retained cortical plasticity may be an important mechanism behind the better cognitive and perhaps somatic health of subjects with high general cognitive ability.

Overall, slow waves reflect the ability of cortical neuron populations to engage in synchronized activity through synaptic connections, and it is affected by ageing, resulting in worse cognitive functioning. This renders slow wave activity a lucrative marker of age-related cognitive problems.

Slow waves have multiple characteristic which reflect different properties of the underlying synchronized neuronal firing. The amplitude of waves reflects the maximum extent of firing synchrony (Vyazovskiy et al., 2008; Nir et al., 2011), the steepness of slopes reflect the rapidness of the buildup and cessation of neuronal firing (Esser et al., 2007; Riedner et al., 2007), frequency – not independently from the two former – reflects the overall synaptic load (Carrier et al., 2011). It has been demonstrated that different slow wave parameters are differently affected by ageing (Carrier et al., 2011), and age-related structural changes in the brain have different effects on density and amplitude (Dube et al., 2015).

While providing more temporal resolution than power spectrum density or the total number of slow waves, SW slope and amplitude are crude approximations of the true morphology of slow waves and therefore can only approximately track the age-related changes of the latter. Due to their highly stereotypical morphology, it is possible to investigate slow waves with a methodology similar to what is used in case of event-related potentials (ERPs) (Key et al., 2005). ERP analysis focuses on the average of EEG signals following several functionally identical events (usually the repeated presentation of a stimulus) and it identifies stereotypical patterns in EEG activity which may be obscured in individual cases but appear in the averaged recording. In the sleep EEG, such analyses were previously performed on a specific type of slow wave, the evoked K-complex (Crowley et al., 2002). K-complexes are morphologically similar to ordinary slow waves which can also appear spontaneously but can also be elicited by stimulation (Halász, 2005). Evoked K-complex components, computed in an identical manner to ERPs in wakefulness, are not only sensitive to the modality of the stimulus with which they were elicited, but also to sex, age and disease (Crowley et al., 2002; Colrain et al., 2010; Colrain and Baker, 2011; Colrain et al., 2011). It is reasonable to assume that the shape of spontaneously occurring NREM slow wave is also subject to such age-related variation but this has never been empirically investigated.

An ERP-like analysis on spontaneous slow waves is possible even in the absence of a trigger stimulus by aligning the EEG signal to a stereotypical morphological element of SWs (such as the negative peak) and averaging accordingly. This analysis reveals not only the major microstructural parameters of slow waves (such as half-wave frequency or ascending/descending slope steepness) but also identifies specific wave components and their precise dynamics at a temporal resolution equal to the sampling frequency of the recording, resulting in substantial additional information about slow wave dynamics.

The goal of this study was to investigate the age-related changes in the high-resolution temporal morphology of spontaneously occurring NREM slow waves using a large existing sleep EEG database of healthy subjects of various ages. As a result of this study we were able to pinpoint candidate electrophysiological markers of healthy ageing beyond the change of major slow wave elements such as amplitude or density. It has been revealed that younger subjects are characterized by faster SW phase transitions both preceding and following negative peaks, suggesting a more rapid buildup and cessation of SW downstates.

## Methods

### Participants

Pre-existing data from 159 healthy subjects (mean age 29.6 years, SD 10.75 years, range 17-69 years; 86 males) from a multi-center database of the Max Planck Institute of Psychiatry (Munich, Germany), the Psychophysiology and Chronobiology Research Group of Semmelweis University (Budapest, Hungary) and the Budapest University of Technology and Economics (Ujma et al., 2014; Pótári et al., 2017) was used in this retrospective study. Data from one 19 year old male subject was excluded due to abnormal average slow wave morphology. Study procedures were approved by the ethical boards of Semmelweis University, the Medical Faculty of the Ludwig Maximilian University or the Budapest University of Technology and Economics. All subjects were volunteers who gave informed consent in line with the Declaration of Helsinki. According to semi-structured interviews with experienced psychiatrists or psychologists, all subjects were healthy, had no history of neurologic or psychiatric disease, and were free of any current drug effects, excluding contraceptives in females. Consumption of small habitual doses of caffeine (maximum two cups of coffee until noon), but no alcohol, was allowed. Six male and two female subjects were light-to-moderate smokers (self-reported), and the rest of the subjects were non-smokers. Further details about participant selection criteria and study protocols can be found in the studies reference above.

### Electroencephalography

All subjects underwent all-night polysomnography recordings for two nights, and data from the second night was used for all analyses. Scalp EEG electrodes were applied according to the 10-20 system (Jasper, 1958) and referenced to the mathematically linked earlobes. Impedances were kept at <8kΩ. EEG was sampled at 250 Hz for 115 subjects, 249 Hz for 29 subjects and 1024 Hz for 15 subjects, always resampled at 250 Hz (see *Slow wave detection*). Sleep EEG was visually scored on a 20 second basis according to standard criteria (Iber et al., 2007). A visual scoring of artifacts was also performed on a 4 second basis. EEG preprocessing was implemented in Fercio’s EEG (©Ferenc Gombos, Budapest, Hungary).

### Slow wave detection

The signal of the electrodes F3 and F4 was band-pass filtered to 0.5-4 Hz (two-way FIR filter implemented in EEGLAB) and averaged. Slow waves were detected on this averaged signal. A slow wave was detected if a negative deflection of the signal persisted for 0.2-1 second and exceeded an amplitude threshold based on previous studies. In order to test the robustness of results across several possible amplitude thresholds, detections were performed using three different values and used in subsequent analyses:

1. Maximal negative deflection <-30 μV, with the lower 50% of the amplitude distribution discarded as putative false positives (Bersagliere and Achermann, 2010) - note that in this case the exact amplitude and distribution thresholds were not identical to the cited study due to the differences in study goals
2. Maximal negative deflection <-37.5 μV, with peak-to-peak amplitude to the subsequent positive deflection >75 μV (Piantoni et al., 2013)
3. Maximal negative deflection <-75 μV, with peak-to-peak amplitude to the subsequent positive deflection >140 μV (Massimini et al., 2004)

Putative slow waves which were detected even partially outside N2 or SWS and/or coincided with a visually detected artifact epoch were discarded. All slow waves detected using each detection threshold were extracted from the EEG signal used for their detection and averaged for each subject, yielding three individual average slow waves for each subject, one according to each detection criterion. In order to control for inter-individual differences in baseline EEG voltage due to sex and body size effects, we also calculated a standardized slow wave for each subject, in which the mean amplitude of the average slow wave was 0 and its standard deviation 1, regardless of raw voltage. Since this was performed for the average waves detected using all three methods, it resulted in a total of six average slow waves for subject, three consisting of EEG voltage time series and another three of standardized values of the former.

Macroscopic features of slow wave morphology frequently used in EEG literature (henceforth referred to as “gross parameters”) were represented by the number of slow waves, their average amplitude (negative peak voltage in μV) and their average slopes – descending downstate (DDS), ascending downstate (ADS), ascending upstate (AUS), descending upstate (DUS), in all cases expressed in μV/second – which were calculated for each subject. In addition to these, for the 30 μV model which used a dynamic detection threshold corresponding to the 50^th^ percentile of the distribution of these relatively small slow waves the actual value of the 50^th^ percentile was also used as a gross parameter.

The fine morphology of slow waves was represented by 501 equally spaced time points including and sampled ±1 second around the negative peak of the average individual slow waves, thus separated by 4 milliseconds. Amplitude values at these time points were obtained using linear interpolation for the subjects whose original EEG sampling frequency was not 250 Hz. The fine morphology of slow waves was calculated using all three detection criteria and both standardized and non-standardized values to test the robustness of results.

Slow wave detection and further statistical analyses were implemented in MATLAB R2017a (The MathWorks, Natick, MA) using EEGLab (Delorme and Makeig, 2004) and custom scripts.

### Statistical analysis

This study aimed to reveal whether the fine morphology of slow waves is affected by ageing beyond its effects on gross parameters: formally, whether the slow wave voltage values included in the measurement of fine morphology account for additional variance in age beyond what is accounted for by gross parameters. Standard least square regression modeling of this relationship was precluded by the large number of predictors: while age and EEG data was only available for 159 subjects, gross parameters and fine morphology of slow waves was represented by 507 predictors (508 in case of 30 μV models), leading to a large degree of model underdetermination. Since fine morphology was expected to be redundant, with highly correlated voltage values across sampling points (see also **Figure 1**), we aimed to solve the problem of underdetermination using least absolute shrinkage and selection operator (LASSO) regression (Tibshirani, 1996). LASSO was implemented in MATLAB with 5-fold crossvalidation. LASSO is an iterative learning algorithm which aims to minimize the following function:

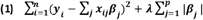

**Figure 1.**
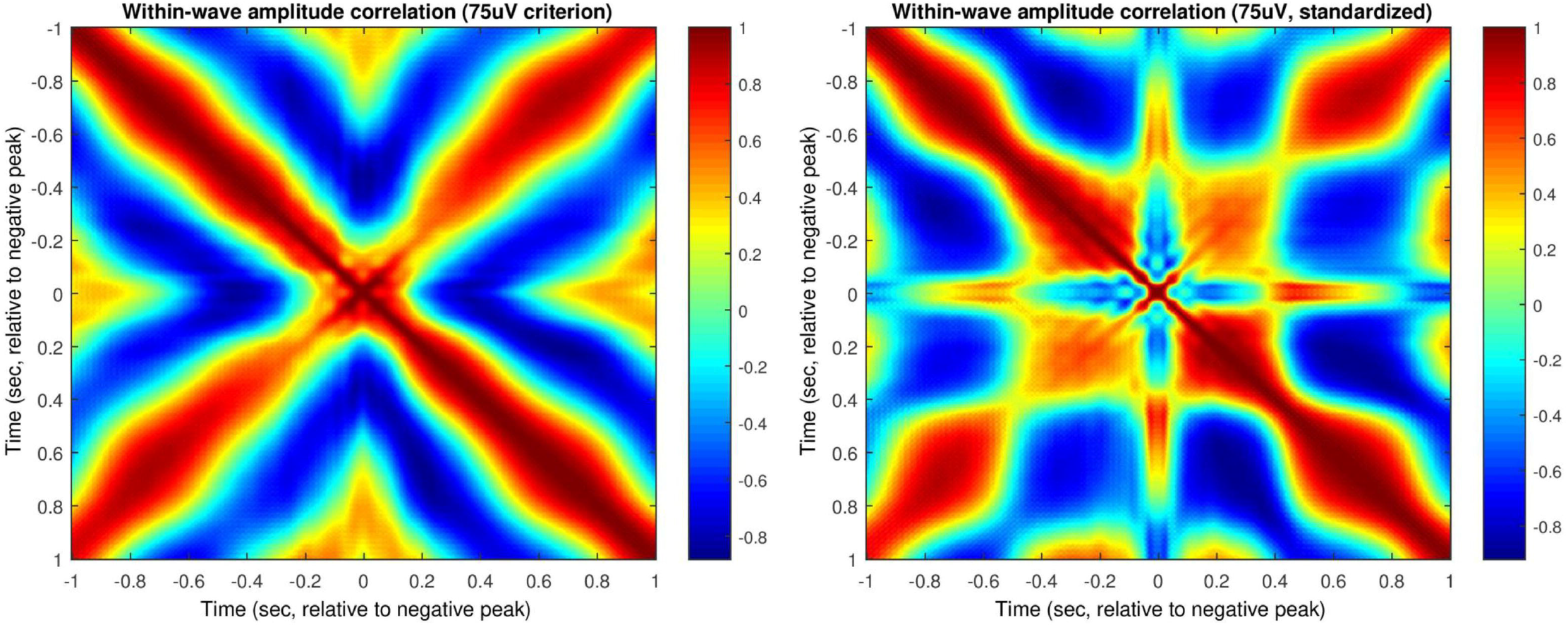
Pearson’s correlation coefficients between slow wave amplitude at 501 equally spaced time points ±1 second around the negative peaks for non-standardized (left panel) and standardized (right panel) slow waves. Note the high positive correlations around the diagonals, representing the similar amplitude of neighboring data points, and the negative correlations between amplitude at time 0 and times ~-0.4 sec and 0.4 sec, indicating that larger negative deflections are preceded and followed by larger positive deflections. The grand average 75 μV slow wave is shown on both subplots for guidance.

, where y_i_ is the dependent variable with n observations, x_ij_ are predictors (total number: p), β_j_ are regression coefficients assigned to the predictors, and λ is a penalty parameter which increases in each iteration of LASSO. It is easy to see that the parts of (1) before λ refer to the residual or prediction error of the regression model: however, in each iteration an additional error term is added for each non-zero regression coefficient |β_j_|, and this additional error is larger for larger values of λ. In other words, with higher values of λ the model is forced to constrain more regression coefficients to 0, leaving only the strongest predictors in the model, until ultimately no predictors remain. LASSO balances the number of predictors against prediction accuracy, finding a value of λ at which enough predictors are left in the model to account for maximal variance but not so many that it would result in underdetermination. Formally, LASSO seeks a λ value for which optimal fit to (1) is achieved in a holdout sample. LASSO is useful when only a subset of the measured predictors are really associated with the dependent variable (Lello et al., 2017) or when a small subset or a single best candidate must be selected from multiple highly correlated predictors (Krapohl et al., 2017) and in the biological sciences it is especially common in animal breeding genetics (de los Campos et al., 2013).

In short, LASSO is able to reliably select few predictors from a large sample of candidates which account for maximal variance in the dependent variable.

## Results

First, the correlation between slow wave amplitude (voltage) values across time points was estimated. This was calculated as the correlation matrix between the voltage values of the individual average slow waves (see Methods) ±1 second before and after the detected negative peaks. Substantial correlation between slow wave amplitude values at different time points would indicate that these amplitude values are redundant and a LASSO regression selecting a smaller subset of independent predictors from them to predict age is appropriate. As expected, slow wave amplitude was strongly correlated across time points, yielding the strongest positive correlations between neighboring time points and the strongest negative correlations between positive and negative peaks. The correlation matrix is illustrated on **Figure 1.**

Next, LASSO regression models were run with age as the dependent variable and gross parameters (number of waves, mean amplitude and all four slopes) as well as amplitude values at each of the 501 time points of the individual average slow waves were entered as predictors. This model construction was repeated for each of the three slow wave detection methods (30 μV cutoff with 50% distribution criterion, 75 μV cutoff, and 140 μV cutoff) and for unstandardized and standardized waves separately, yielding a total of six LASSO model runs. **Figure 2** illustrates the change in mean squared error in the holdout sample as a function of λ parameter values in a LASSO model run, while Figure 3 illustrates the change in the number and identity of predictors as a function of λ.

**Figure 2.**
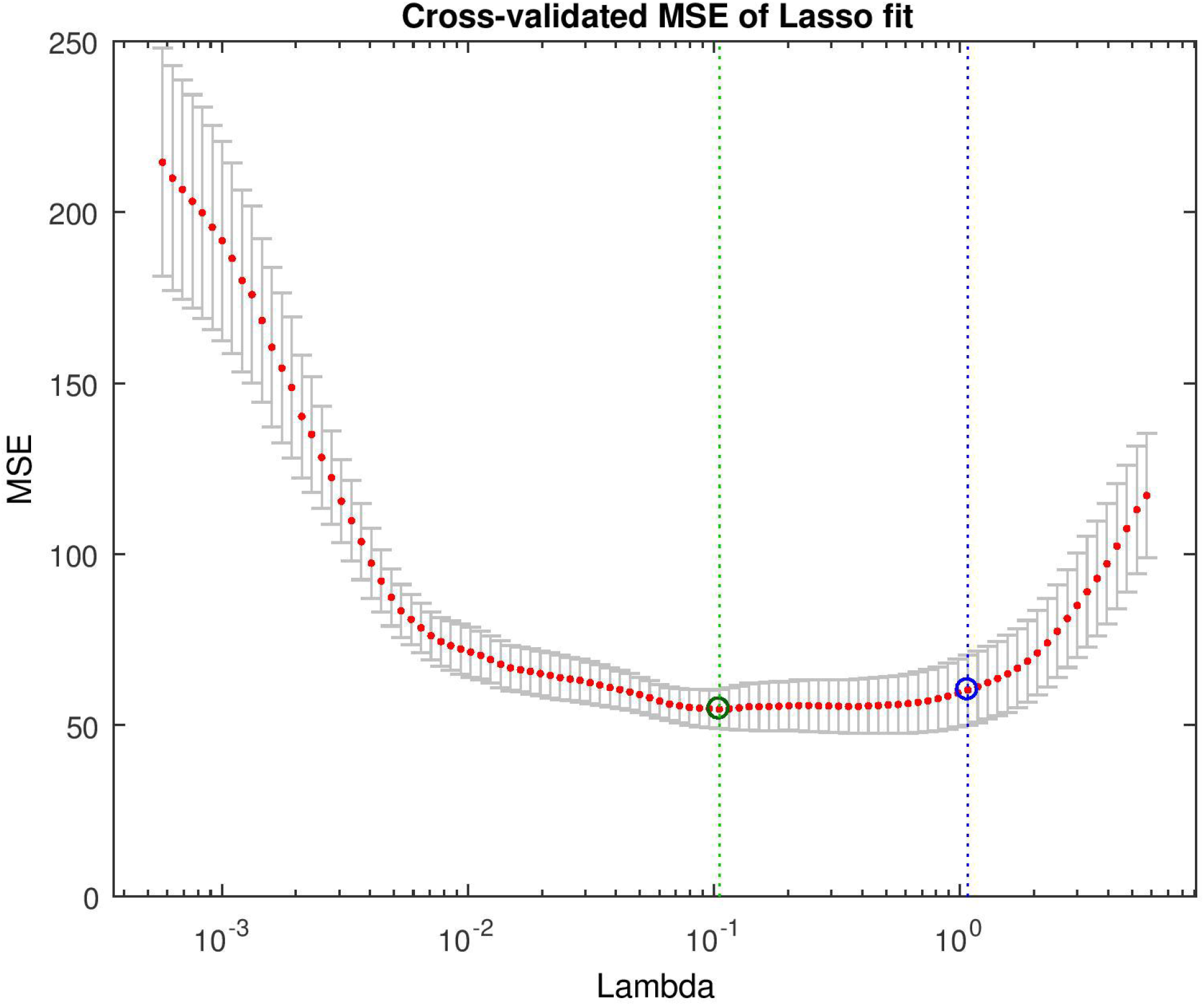
LASSO model run for 75 μV unstandardized slow waves with the MATLAB built-in lassoplot() function. With the initial low values of λ, the number of predictors in the model and consequently the degrees of freedom is high, resulting in high mean squared error with very large confidence intervals. As the value of λ increases, only large-effect predictors are left in the model, resulting in more precise estimates. The green dot indicates the λ value at which optimal prediction performance is reached in the holdout sample, while the blue dot indicates the value minimum prediction error plus one standard deviation.

**Figure 3.**
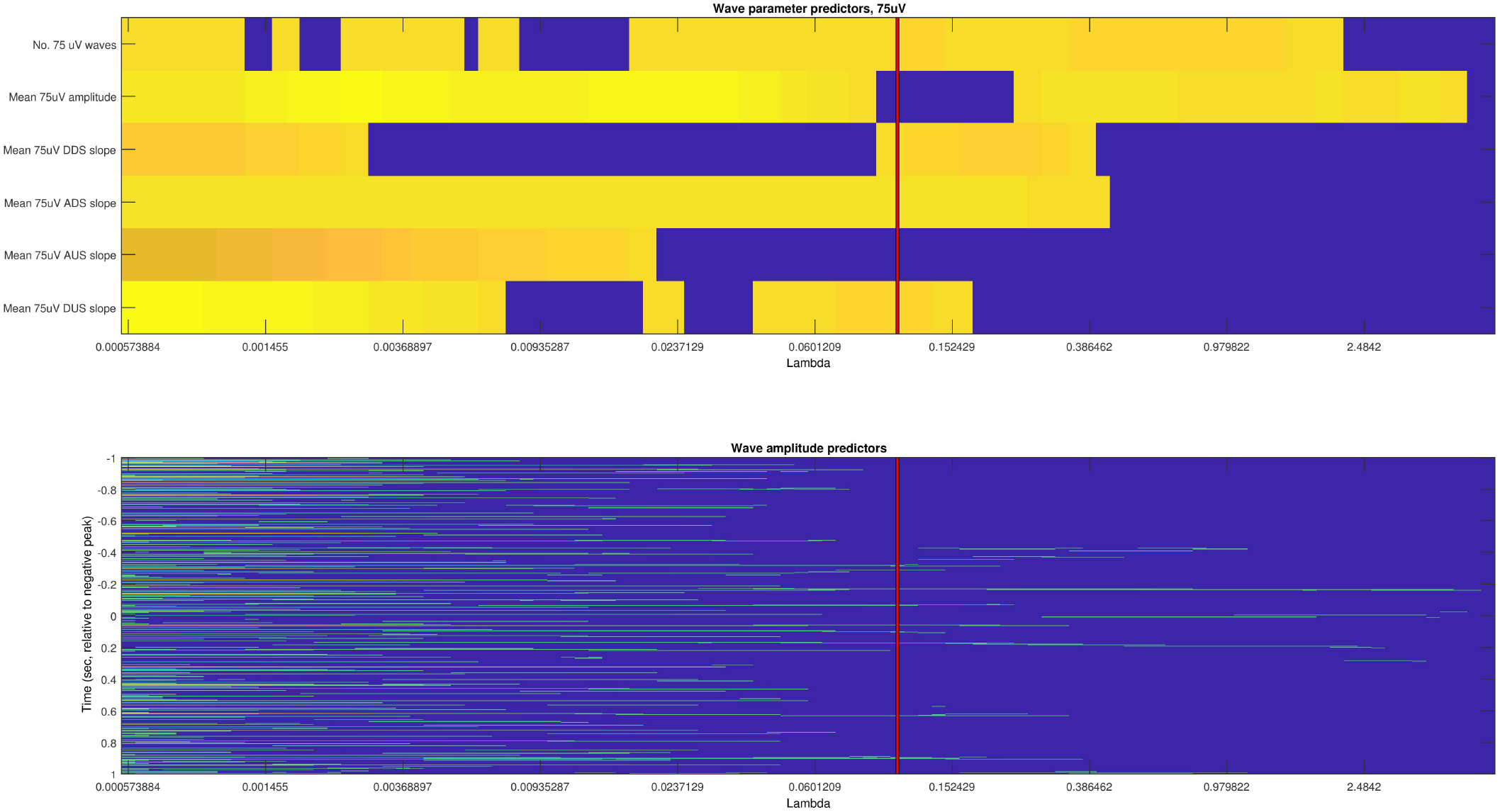
Predictor regression coefficients of the LASSO model built for 75 μV unstandardized slow waves. Gross slow wave parameters (number of waves, mean amplitude and all four slopes) and amplitude values at 501 equally spaced time points ±1 second around the negative peaks of average slow waves are shown separately on the upper and lower panels, respectively, but were always entered in the model together. Color (on an arbitrary scale due to range differences, with darker yellow indicating higher regression coefficients) indicates the significance of the parameter indicated on axis y at the λ value indicated on axis x. Dark blue areas indicate that the regression coefficient for the predictor at the given λ value was set to zero, indicating no advantage to the inclusion of the predictor over the cost of its increasing degrees of freedom. The red vertical line is identical to the green dot of Figure 2 and indicates the λ value at which the optimal tradeoff between model complexity and prediction performance was reached. The fact that this line crosses many significant predictors among the amplitude values (lower panel) illustrates that wave point amplitudes contributed to the prediction of age over the effect of gross parameters.

According to the LASSO model results, amplitude at certain slow wave time points remained as significant predictors in the best-fit model in case of all six slow wave detection methods. In other words, the inclusion of slow wave amplitude at specific time points always resulted in a significantly more accurate prediction of age than the inclusion of gross parameters only (**Figure 3**). That is, age-related changes in slow wave morphology were not limited to a change in the number, amplitude or overall slope of these waves. **Figure 4** illustrates which specific time points of individual average slow waves remained significantly predictive of age beyond the effects of gross parameters.

**Figure 4.**
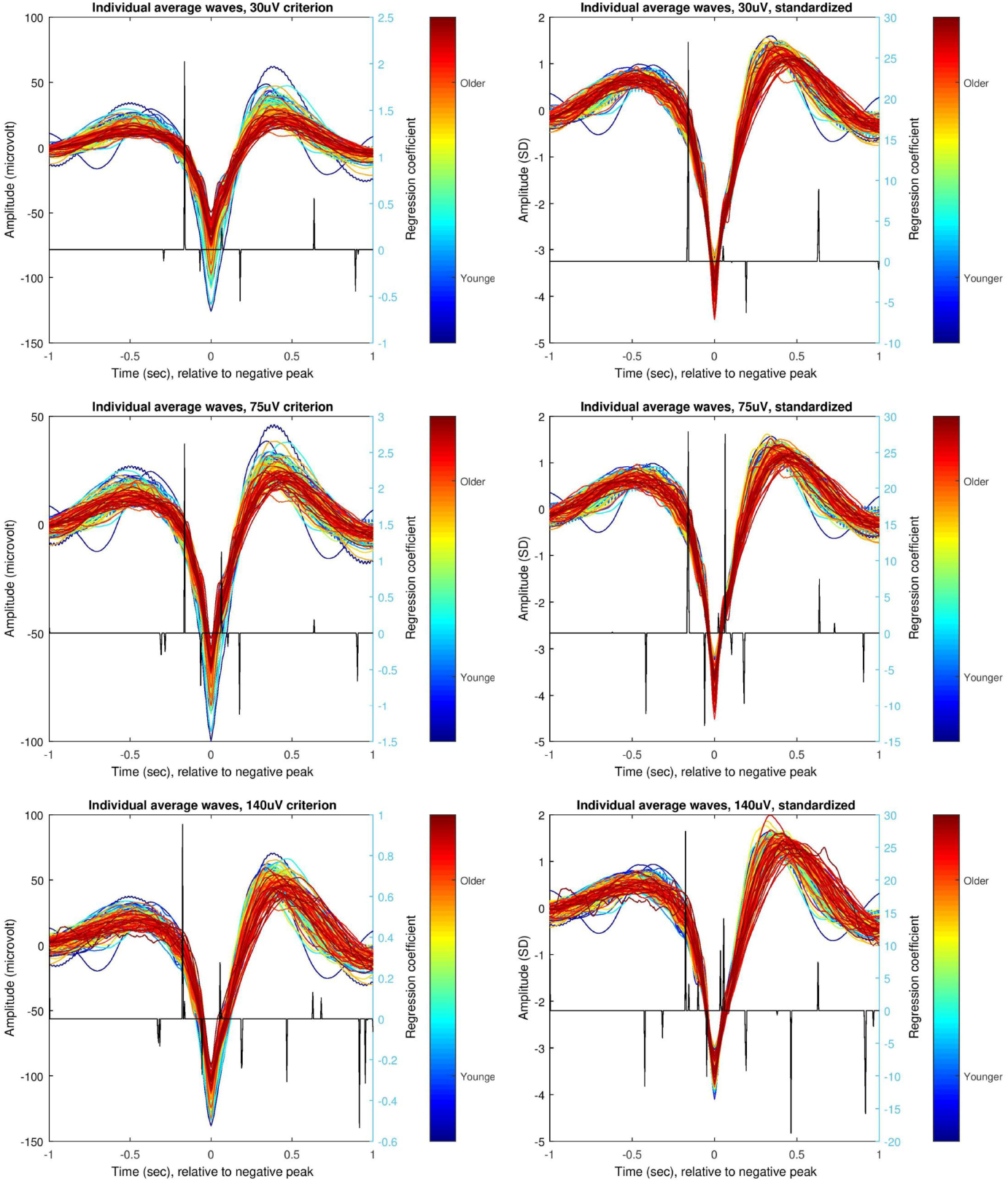
Uncorrected individual average standardized (left panels) and unstandardized (right panels) slow waves, triggered to the negative peak. Data is shown for each of the three amplitude criteria used in robustness analyses. Each subjects’ individual average slow wave is shown overlapping to the others, and color coded according to the rank order of the subject’s age. Note a strong age effect on slow wave negative peak amplitude in unstandardized but not in standardized waves. A single black line indicates LASSO regression coefficients in the best-fit model (indicated by a green dot on Figure 2 and a red line on Figure 3) at the corresponding slow wave time point, and it is expressed as years/amplitude standard deviation (standardized waves) or years/μV (unstandardized waves), in all cases corrected for the effects of gross parameters. The sign of the regression coefficient indicates the accurate direction of the association between slow wave amplitude and age.

The incremental validity of the statistical model predicting subject age from slow wave morphology in addition to gross parameters is showcased by the higher correlations between predicted and observed values with this method (**Figure 5**). When a prediction of age is performed based on the regression coefficients of unstandardized 75 μV gross parameters in the best-fit LASSO model, the correlation between predicted and observed age values is 0.61, while with the inclusion of wave morphology effects this correlation increases to 0.81 (a significant difference with z=3.69, p=0.0002 based on Fisher’s r-to-z transformation).

**Figure 5.**
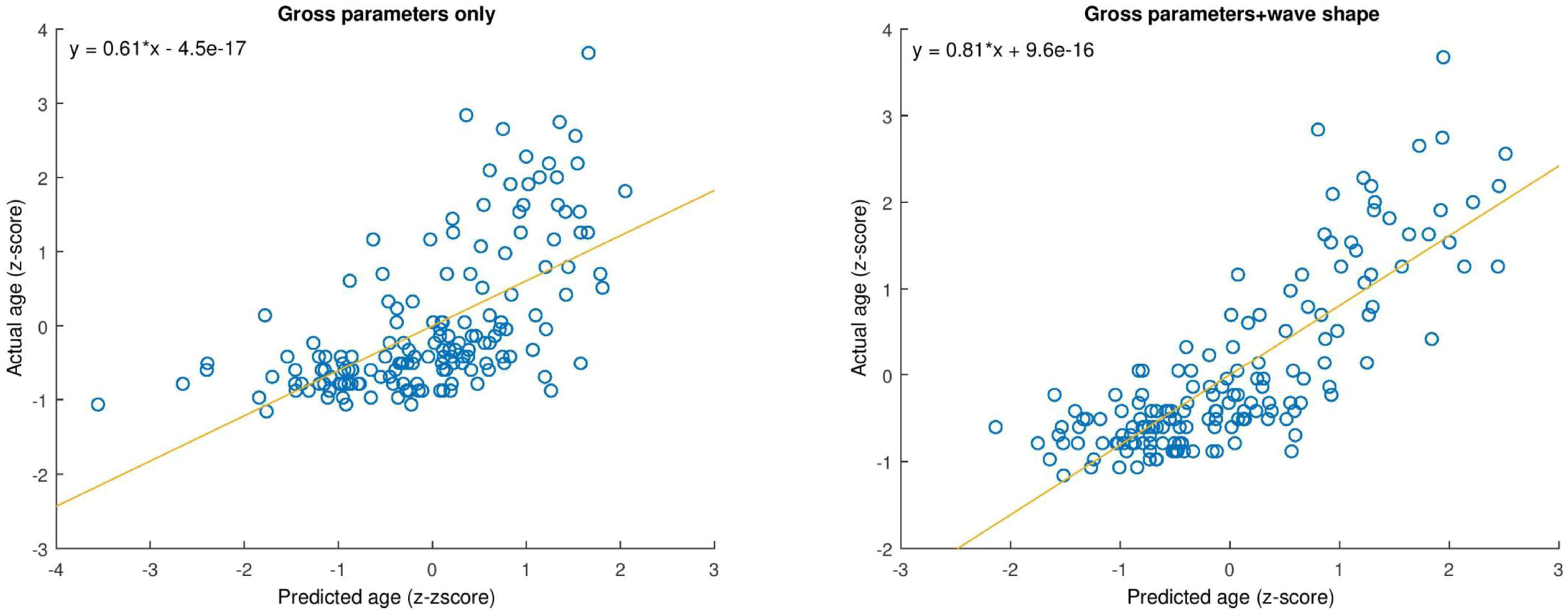
The correlation between predicted and observed age using gross wave parameters only (left panel) or individual slow wave morphology in addition to gross parameters (right panel). Both predicted and observed age is standardized so that the regression coefficient is equal to Pearson’s correlation coefficient.

Last, we investigated the overall temporal dynamics of the association between ageing and fine slow wave morphology: that is, we created summary statistics about which parts of slow waves contained amplitude values which contributed additional explained variance towards age beyond the effect of gross parameters. We created 16 temporal bins (each of a length of 125 msec) of the 501 amplitude values and summed the number of non-zero coefficients in the best-fit LASSO models based on all six slow wave detection methods in each of the bins. The results are illustrated on **Figure 6.** Slow wave morphology was most frequently associated with age in the descending downstates preceding and ascending downstates immediately following the negative peak, as well as during descending upstates following the slow wave peak by almost 1 second. Downstates were especially enriched for such amplitude values, but their distribution on very specific parts of wave slopes may explain why slope steepness failed to fully account for the covariance of slow wave morphology and age.

**Figure 6.**
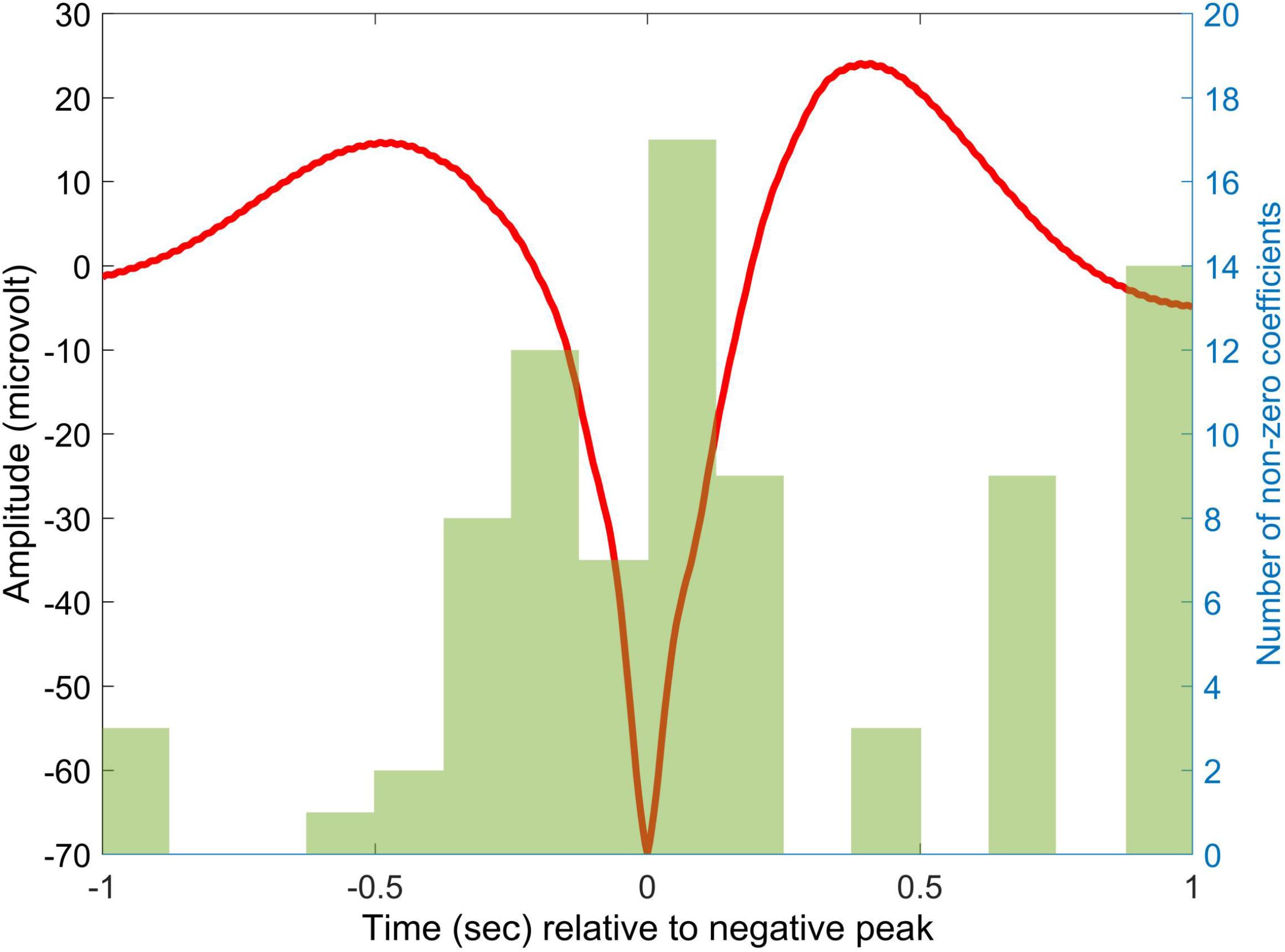
The number of non-zero coefficients in the best-fit LASSO models, summed across all six models based on different slow wave detection methods and for 125 msec time bins. The grand average slow wave based on the averages of 75 μV slow waves is shown for guidance.

In sum, a penalized regression model demonstrates that the slow wave gross parameters (number of waves, amplitude and slope) regularly used by EEG studies of NREM sleep and ageing capture only part of the age-related change of these waveforms, and the fine morphology is more specifically affected by ageing.

## Discussion

This study investigated age-related changes in the high-resolution morphology of slow waves in a large sample of subjects. It replicated previous findings (Carrier et al., 2011) about an age-related reduction in slow wave number, amplitude and slope even in this sample of mostly young and middle-aged subjects. It was found, however, that age-related changes in the number of waves, amplitude and slope fail to fully account for the alterations in slow wave morphology. Even after correcting for the effects of these predictors in a penalized regression model, fine morphology added significant additional variance towards age. Notably, the correlation between gross parameters and age was 0.6, compared to the previously reported 0.49 for slow wave density in 82 subjects (Carrier et al., 2011) or 0.28 for log delta power in 211 subjects (Schwarz et al., 2017)^1^, and 0.28 and 0.62 for relative delta power in two subsamples (N=87 and N=72, respectively) of the present study (Pótári et al., 2017) (The sign of the correlation coefficient was ignored in all cases). The correlation between a predicted age based on both gross parameters and fine morphology was 0.8 with actual age, which is higher than what was obtained in previous large-sample studies, highlighting the importance of age-related changes in the fine morphology of slow waves.

Slow wave morphology was generally associated with age during phase transitions, especially late descending and early ascending downstates and late descending upstates (possibly indicating the initiation of the next downstate). Notably, amplitude around slow wave negative peaks was less frequently associated with age, suggesting that slow wave amplitude as a gross parameter adequately captured the association between this aspect of slow wave morphology and age. These findings suggest that age-related changes in slow wave morphology especially affect the speed and degree of the initiation and cessation of slow wave up- and downstates.

These findings are interpretable in light of previous literature using high density scalp EEG, coregistered invasive and scalp EEG in humans or computer and rodent models. Slow waves accurately reflect the onset and cessation cortical neuronal firing, even in human recordings (Csercsa et al., 2010; Nir et al., 2011). Consequently, the shape of the slow wave reflects how rapidly and to what extent neuronal firing is synchronized (Esser et al., 2007; Riedner et al., 2007; Vyazovskiy et al., 2007). In line with the view that slow waves reflect a homeostatic response to long-term potentiation taking place in the preceding wakefulness (Tononi and Cirelli, 2014), frequent, high-amplitude slow waves with steep slopes are seen under conditions of high homeostatic pressure for sleep, such as early in the night (Esser et al., 2007; Riedner et al., 2007) or in case of young subjects with a higher degree of neuronal plasticity (Feinberg and Campbell, 2013). Based on these studies, the current results can be interpreted as a more rapid initiation and cessation of synchronized cortical neuronal firing in younger subjects, in line with the finding that in older subjects, both the homeostatic need for slow waves and the ability to respond to this need by slow wave generation is reduced (Mander et al., 2017).

While the previously reviewed literature suggests that the amplitude, the slope and to a degree even the number of slow waves is mainly a function of synaptic plasticity, these slow wave gross parameters are not redundant, evidenced by an imperfect within-wave correlation between slopes, amplitude and frequency. While the literature on this topic is scarce, (Botella-Soler et al., 2012) found a low correlation between descending and ascending slope steepness in corticographically recorded slow waves. In the present study, within-subject correlations between average gross parameters for the representative case of 75 μV waves averaged 0.6 (SD=0.47), with the lowest correlation between ascending downstate slopes and other gross parameters. A principal component analysis extracted a first unrotated factor accounting for 65% of the variance of all gross parameters, showing substantial but not nearly perfect interdependence.

In line with the hypothesis that slow wave parameters are relatively independent, their age-related changes also show different patterns. Age-related changes in slow wave slopes remained significant after controlling for the change in amplitude (Carrier et al., 2011). A structural equation modeling study (Dube et al., 2015) combining imaging with polysomnography revealed that age-related reductions in slow wave density and amplitude are almost completely accounted for by cortical thinning. However, different structural imaging changes accounted for the age-related reduction of density and amplitude, highlighting that these measures rely on different anatomical structures.

Age-related changes in any slow wave measure are important because alterations in NREM sleep function are thought to play a functional role in age-related cognitive decline (Mander et al., 2017). Following up animal experiments demonstrating the effect of sleep deprivation on β amyloid accumulation (for a summary, see (Ju et al., 2013)), structural equation modeling has demonstrated that the relationship between β amyloid accumulation and reduced memory function is mediated by slow wave activity (Mander et al., 2015). This suggests that an important part of Alzheimer pathology may be related not directly to the β amyloid burden, but to the resulting disruption of NREM sleep (Lucey and Holtzman, 2015). While it would be tentative to see both the cognitive symptoms and NREM sleep changes in Alzheimer’s disease as an extreme case of normal aging, it must be considered that while normal ageing involves a reduction of a wide range of slow frequency activity not limited to delta, very low frequencies are especially affected in Alzheimer’s disease (Mander et al., 2017). The molecular composition of β amyloid proteins is also different in normal ageing and Alzheimer’s disease (Piccini et al., 2005). Therefore, it requires further research to establish the degree to which slow wave changes in normal ageing and in dementia are analogous.

While the translational potential of slow wave research to clinical practice is limited at the moment, some evidence suggests that cognitive functioning in wakefulness can be improved by artificially increasing slow wave activity in NREM sleep through heating, auditory or electrical stimulation, including in old age (Ladenbauer et al., 2017; Papalambros et al., 2017; Wilckens et al., 2018). Therefore, the significance of the present study is to demonstrate that age-related changes in slow wave morphology possibly underlying the age-related reduction of cognitive function are specific, not fully captured by gross parameters and they especially include the rapidness of phase transitions, in line with previous findings about the additional predictive value of slopes over amplitude towards age (Carrier et al., 2011). Methods able to restore this specific property of slow waves in high age or dementia may be especially effective in improving cognitive functioning.

## Footnotes

1 For Carrier et al 2011, the correlation was calculated from F=13.9 (Fig. 2) for the dichotomous age difference, corresponding to d=0.835 and r=0.39 at N=82. This correlation was corrected for the downward bias of dichotomization assuming a cutoff at 52.4% of the sample based on subsample sizes, corresponding to the normal deviate z=0.06 and the normal ordinate 0.4. For Schwarz et al 2017, the correlation was calculated from the t-values in Table 4.

